# Deep Learning Modelling of Androgen Receptor Responses to Prostate Cancer Therapies

**DOI:** 10.1101/2020.01.15.908384

**Authors:** Oliver Snow, Nada Lallous, Martin Ester, Artem Cherkasov

## Abstract

Gain-of-function mutations in human Androgen Receptor (AR) are amongst major causes of drug resistance in prostate cancer (PCa). Identifying mutations that cause resistant phenotype is of critical importance for guiding treatment protocols as well as for designing drugs that do not elicit adverse responses. However, experimental characterization of these mutations is time consuming and costly, and therefore predictive models are needed to anticipate resistant mutations and to guide drug discovery process. In this work, we leverage experimental data collected on 69 clinically observed and/or literature described AR mutants to train a deep neural network (DNN) to predict their responses to currently used and experimental AR anti-androgens. We demonstrate that the use of DNN provides more accurate prediction of the biological outcome (inhibition, activation, no-response) in AR mutant-drug pairs compared to other machine learning approaches and also allows the use of more general 2D descriptors. Finally, the developed approach was used to predict the effect of the latest AR inhibitor darolutamide on all reported AR mutants.

## Introduction

Resistance to drug treatments is a common occurrence across many diseases but it is especially prevalent and lethal in cancers, where it represents a major obstacle for long term therapies. The acquired drug resistance can be caused by a number of different mechanisms by which cancer cells can escape the treatment.^1^ One of the major ways is the development of gain-of-function mutations where the target protein becomes altered under selective therapeutic pressure, rendering the drug ineffective. Gain of function mutations are of particular importance in prostate cancer (PCa) where resistance is a common and often deadly occurrence.^2^

The main drug target in PCa is the androgen receptor (AR), a nuclear hormone receptor whose increased activation is one of the principal drivers of PCa. Multiple decades of research on the AR has led to a number of targeted drug treatments that have significantly improved patient survival and well-being. However, despite the major gains in AR targeted treatments, resistance invariably develops to all current drugs.^3,4^ One important aspect of resistant AR mutants is that they do not simply render the drug ineffective but can even turn the drug from an antagonist into an agonist, thus promoting cancer growth. This characteristic seems to be unique to the AR and thus emphasises the need to identify gain-of-function mutations that cause this phenotype so that patients can be screened and taken off treatment before resistance develops. ^5^ Additionally, understanding and predicting those mutations that cause resistance will enable us to design better drugs that might avoid this mechanism in the future.

Previous research had identified a number of new AR mutants from patient DNA using next generation sequencing technology.^6^ The activation of these mutants were then measured (along with others documented in the literature) in response to increasing doses of the current and upcoming AR drugs were then experimentally confirmed in PCa cell lines. Building upon the previous work, additional mutants listed in the McGill Androgen Receptor Gene Mutations Database^7^ were screened against the same drugs, resulting in a high quality dataset of approximately 50 AR mutants tested against 7 anti-androgen drugs.

Naman et al.,^8^ in 2016 introduced QSAR (quantitative structure-activity relationship) modelling as a binary classification of the AR mutant responses to common PCa drugs and endogenous human steroids using structure-based 4D QSAR descriptors.^9–12^ The QSAR model was able to identify a *de novo* AR mutant that showed a resistant phenotype to the current anti-androgen drugs. However, the approach was relatively simplistic and despite achieving an accuracy of ∼80%, the model generated several false positive predictions for the external validation set. Importantly, the use of binary antagonist/agonist classification reduced the complexity of the AR mutant responses that can range from full antagonism to full agonism and also includes partial agonistic response and no-response (non-functional AR mutants) categories.

Recent advances in a machine learning theory have led to significant progress and qualitative changes in QSAR and chemoinformatics practice. Interpretable linear QSAR models become increasingly replaced by support vector machines (SVM), random forest (RF), artificial neural networks (ANN) and other non-linear techniques that demonstrate robust performance on large biological datasets. ^13,14^ More recently the progress in deep learning techniques along with the increasing availability of biological data has brought DNNs into the drug modelling spotlight.^15^ Deep networks have been particularly effective due to their ability to learn more complex non-linear trends from larger datasets and their lower requirements for input representations, i.e. lesser need for precise descriptor engineering. ^16,17^

In this work, we utilized a deep neural network (DNN), along with general proteochemometric descriptors to predict more differentiated AR mutant-drug responses on significantly extended experimental dataset, compared to our previous study. Furthermore, we implemented a structure-independent protocol, where by using protein sequence-based descriptors and 2D drug fingerprints, we avoided the drug docking step in feature construction, saving time and making the model more generalizable. Notably, such an approximation allows the consideration of mutations that occur outside of the ligand-binding domain (LBD) of the AR, where no structural information is available. As mentioned above, the resulting DNN model distinguishes four AR mutant response phenotypes and provides rather accurate discrimination between them.

## Results and discussion

### Dataset

AR mutant data was collected as described by Lallous et al.,^6^ in which single and multiple amino acid substitutions of the AR were expressed in PC3 cells. Responses to six different anti-androgens (bicalutamide, ^18^ enzalutamide,^19,20^ hydroxyflutamide,^21^ ARN509, ^22^ VPC13566, and VPC135789) were measured in a luciferase-reporter assay. An example of the measured responses can be seen in Figure 1 where 3 of the main phenotypes (agonist, antagonist, and mixed response) are displayed for the mutant W742C. The dose-response curve data consisting of 11 increasing drug concentrations and their transcriptional activity values (normalized to wild-type) were converted to 4 classes corresponding to the above phenotypes with an additional non-responsive phenotype. The dataset was split into a training set, upon which cross validation and hyperparameter tuning was performed, and a test set to evaluate the performance of the methods.

**Figure 1:**
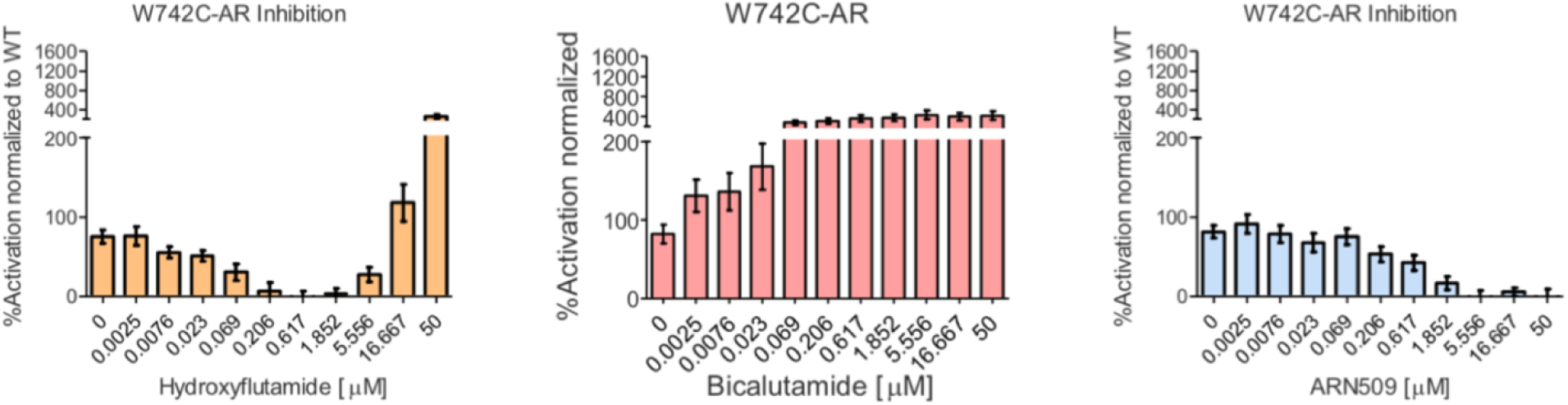
**Example AR mutant drug response phenotypes**

### Feature calculation

Using the fasta-formatted amino acid sequences of the AR variants (trimmed to only include the range in which the current substitutions occur), mutant sequences were generated for each single or multiple mutant in the dataset. These sequences were then used to generate z-scale descriptors using the *iLearn* python package.^23^ Z-scale descriptors characterize each amino acid by 5 physico-chemical values and have been shown to consistently match or outperform other sequence based descriptors on benchmark datasets.^24,25^ Drug molecules were encoded with extended connectivity fingerprint (ECFP; also known as Morgan fingerprint) descriptors using the RDKit python package and a radius of 2, resulting in a 2048 bit vector for each molecule.^26,27^ Z-scale descriptors were then standardized to zero-mean and unit variance while Morgan fingerprints were left as one-hot encoded vectors.

Given that certain drug response phenotypes are less frequent than others (resistant phenotype is relatively rare) there is a high imbalance between classes that can negatively affect the accuracy of the model (the model can get high accuracy by just predicting the majority class). To address this imbalance, we over-sample the minority classes according to the borderline synthetic minority over-sampling technique (SMOTE) method which outperformed the regular SMOTE method and random oversampling in cross validation.^28,29^

### Baseline methods

To justify the use of the more complex DNN method for this task, we benchmarked the corresponding results against outcomes from SVM and RF models. In particular we have obtained 5-fold cross validation predictions by SVM and RF and utilized area under the receiver operating characteristic curve (AUC) statistics. SVM used an radial basis function kernel with *gamma* = 0.001 and the RF was built with *# of estimators* = 50 and *max depth* = 10. Oversampling was performed within the cross validation loop so as not to bleed any information into the validation set.

### DNN model training

The general overview of our approach can be seen in Figure 2, where the network takes in protein sequence and 2D chemical descriptors and outputs 4 classes of dose-response curve. The DNN was also optimized to find the best hyperparameters using 5-fold cross validation. The optimal architecture was found to be two hidden layers with 128 neurons in the first layer and 32 in the second layer, which is a relatively small network but is a suitable amount of weights to train given the small number of training examples. Categorical cross-entropy loss was used, with a batch size of 16 and the ADAM optimizer for training.^30^ Each hidden node had a rectified linear unit (ReLU) ^31^ activation applied and the output layer had a softmax activation applied.^32^ Additionally, to reduce over-fitting of the network, a dropout rate of was applied to each hidden layer and early stopping was used to terminate training if validation loss did not improve over 20 epochs.

**Figure 2:**
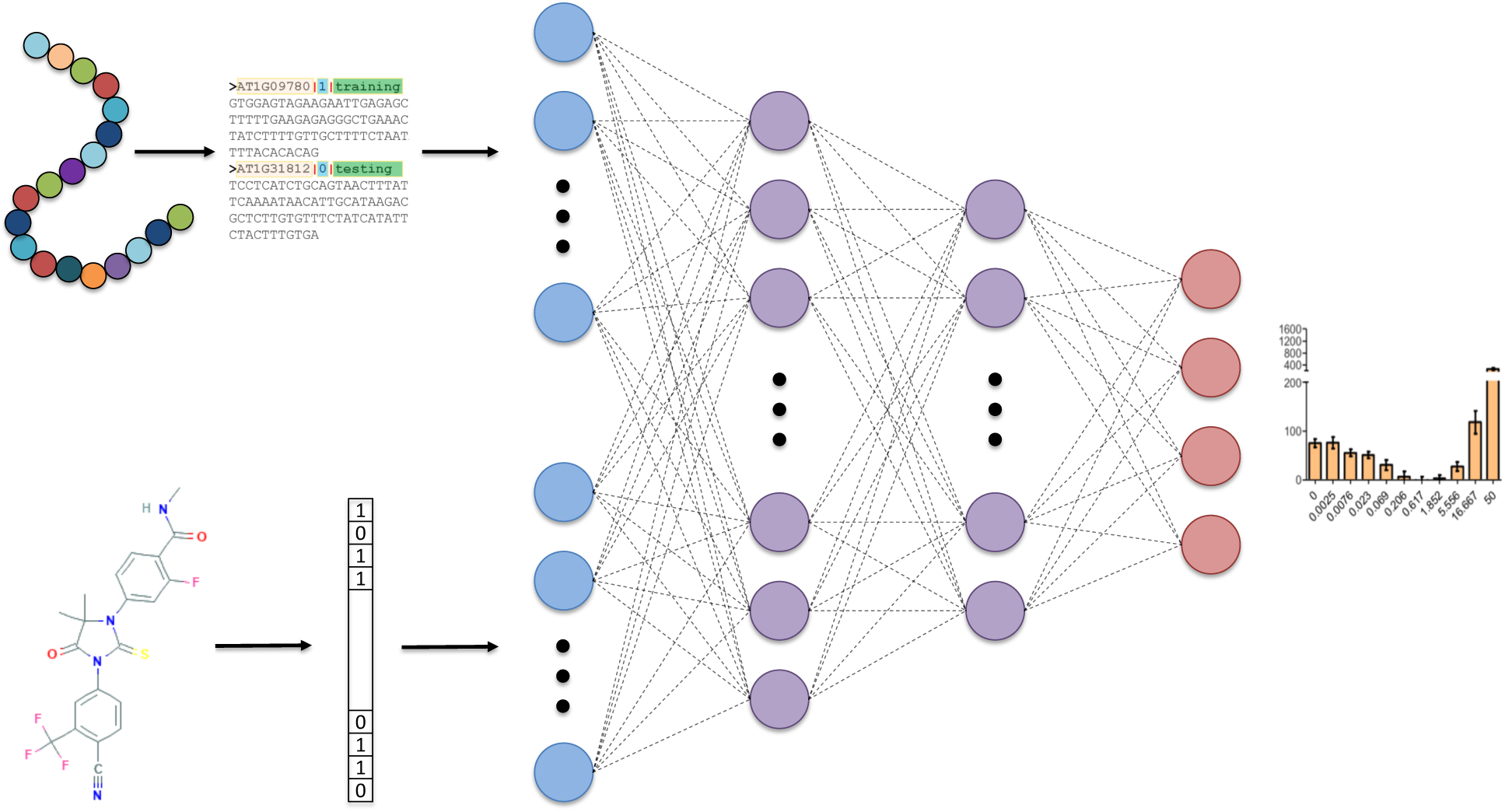
Overview of DNN model taking sequence and 2D chemical descriptors as input, 2 hidden layers with 128 and 32 nodes respectively, and a 4 node output to classify four categories of dose response curves

### Prediction results

Interestingly, SVM and random forest have fairly high AUC on the training set despite the large dimension of the input and the DNN performs only slightly better than SVM and the same as random forest (Figure 3). However, further investigation into precision and recall shows that both baseline methods over-predict for the majority class, a bias that the AUC does not fully capture. This can be seen in Table 1 where the DNN significantly outperforms the baselines in precision, recall and F1 score.

**Table 1:** Classification report for DNN and baselines.

|  | Precision | Recall | F1 Score |
| --- | --- | --- | --- |
| DNN | 0.79 | 0.80 | 0.79 |
| RF | 0.59 | 0.58 | 0.56 |
| SVM | 0.69 | 0.69 | 0.63 |

**Figure 3:**
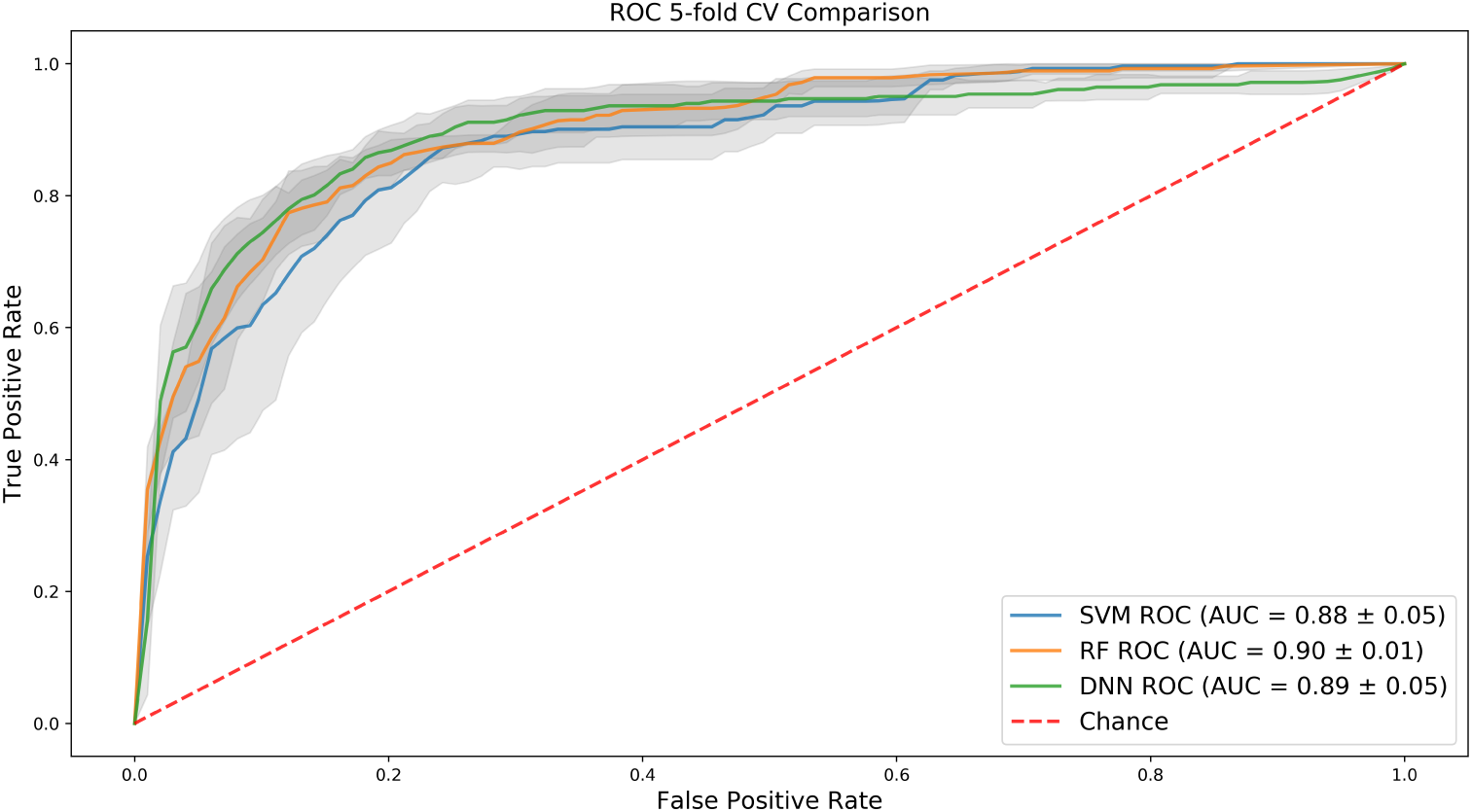
**Mean ROC curves over 5-fold CV for SVM, RF, and DNN with 1 standard deviation shown in grey**

The shallow models had difficulty discriminating between the antagonist and u-shaped phenotypes and thus do not generalize well to validation data. This can be seen clearly in Figure 4 with a comparison of confusion matrices for the three methods on the test set. The DNN predictions are noticeably better than baselines at identifying the non-responsive phenotype and is more sensitive at distinguishing between the mixed (U-shaped) response and antagonist response which are subtly different.

**Figure 4:**
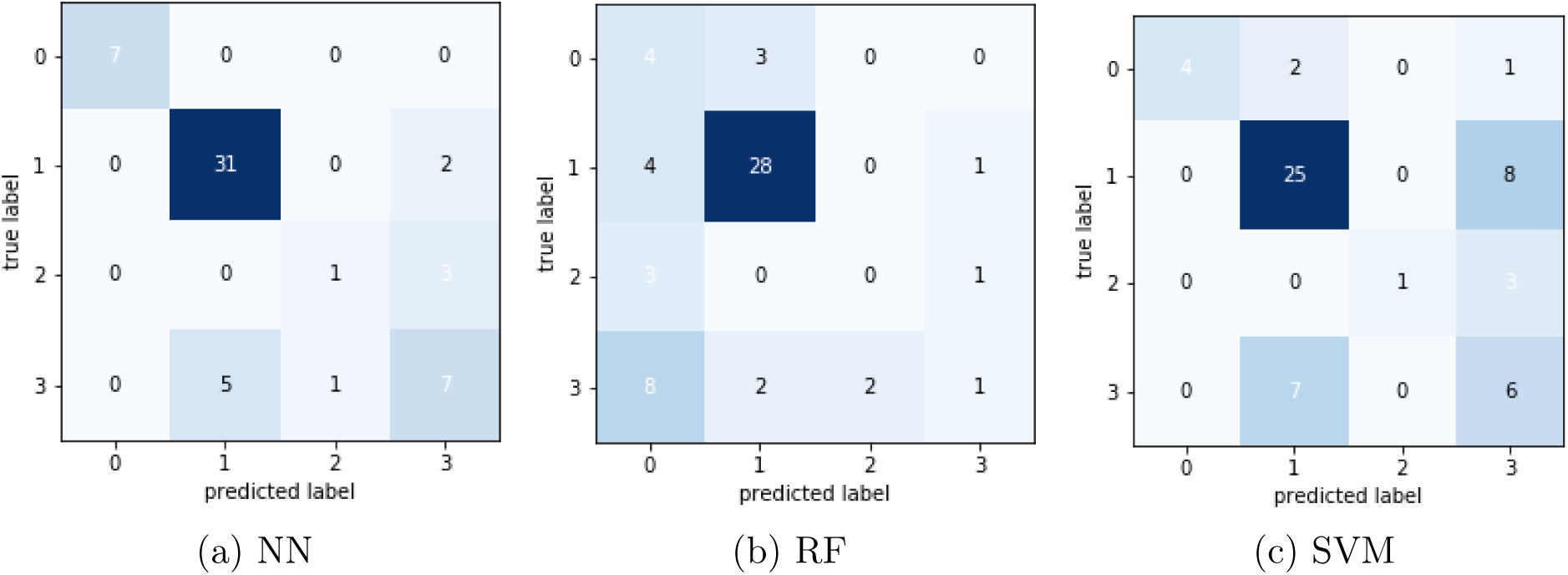
Confusion matrices of three methods on test set;class 0 = non-responsive, class 1 = antagonist, class 2 = agonist, class 3 = mixed response

We next used the trained DNN model to predict the response of 44 common mutants in combination with darolutamide, a leading AR antagonist which has not yet shown a resistant, mixed-response phenotype. To accomplish this, we compute Z-scale descriptors and Morgan circular fingerprints according to the same protocol as above and then use our trained DNN model to predict on these descriptors. Our model predicts that of these 44 mutants, 12 are predicted to be non-responsive, 31 are predicted to be antagonized by the drug, and a single mutant, E666D is predicted to have the resistant (U-shaped) phenotype. Further cell-line experiments should be conducted to confirm these predictions and to investigate whether the predicted resistant mutant might soon be seen in patients with increased treatment with darolutamide.

## Conclusions

In this work we have demonstrated the potential of deep learning in combination with flexible and informative proteochemometric descriptors to predict adverse drug responses of a wide range of acquired AR mutations observed in clinical samples and reported in the literature over the years. As more data becomes available for this problem, the superior accuracy and ease of use of our DNN approach would be expected to grow. This modelling approach could also easily be extended to other protein targets and predict for other drug compounds, either experimental or in clinical use. The developed DNN model would also be of high practical utility, either in the clinical context by warning of resistance causing mutations to current therapies, or in the drug development context to screen lead compounds for their likelihood to elicit resistant phenotypes. Further work could build on this model to take a Bayesian approach in order to give uncertainty of predictions which could then be incorporated in an active learning design to search the vast space of single, double, or multiple mutants that will be most likely to cause resistance to our best AR antagonist therapies. ^33^

